# Powerful Gene Set Analysis in GWAS with the Generalized Berk-Jones Statistic

**DOI:** 10.1101/361436

**Authors:** Ryan Sun, Shirley Hui, Gary D. Bader, Xihong Lin, Peter Kraft

**Affiliations:** Department of Biostatistics, Harvard T.H. Chan School of Public Health, Boston, MA 02215, USA; The Donnelly Center, University of Toronto, Toronto, Ontario M5S 3E1, Canada; Department of Epidemiology, Harvard T.H. Chan School of Public Health, Boston, MA 02215, USA.

## Abstract

A common complementary strategy in Genome-Wide Association Studies (GWAS) is to perform Gene Set Analysis (GSA), which tests for the association between one phenotype of interest and an entire set of Single Nucleotide Polymorphisms (SNPs) residing in selected genes. While there exist many tools for performing GSA, popular methods often include a number of ad-hoc steps that are difficult to justify statistically, provide complicated interpretations based on permutation inference, and demonstrate poor operating characteristics. Additionally, the lack of gold standard gene set lists can produce misleading results and create difficulties in comparing analyses even across the same phenotype. We introduce the Generalized Berk-Jones (GBJ) statistic for GSA, a permutation-free parametric framework that offers asymptotic power guarantees in certain set-based testing settings. To adjust for confounding introduced by different gene set lists, we further develop a GBJ step-down inference technique that can discriminate between gene sets driven to significance by single genes and those demonstrating group-level effects. We compare GBJ to popular alternatives through simulation and re-analysis of summary statistics from a large breast cancer GWAS, and we show how GBJ can increase power by incorporating information from multiple signals in the same gene. In addition, we illustrate how breast cancer pathway analysis can be confounded by the frequency of *FGFR2* in pathway lists. Our approach is further validated on two other datasets of summary statistics generated from GWAS of height and schizophrenia.

## Introduction

A common objective in genetic association studies is to search for associations between phenotypes and genomic constructs that are larger than a Single Nucleotide Polymorphism (SNP). One popular unit of analysis is the set of all SNPs that are located near a list of related genes; inference on these sets is generally referred to as gene set analysis (GSA) or pathway analysis^1^. In recent years, GSA has successfully identified novel gene sets associated with a wide range of outcomes^2–4^.

GSA does not yet possess the popularity of individual SNP approaches^5^ such as the Genome-Wide Association Study (GWAS), but there are advantages to testing for associations at a higher level^6^. Many biological processes are driven by mechanisms involving more than one variant, and thus set-based inference may offer more useful interpretations^7^. In addition, set-based inference can increase power over individual-SNP methods by pooling many weaker pieces of evidence into a larger, more detectable signal^8^. GSA can also improve power by alleviating the multiple testing burden of GWAS^9^.

However, realizing the aforementioned benefits is hampered by a lack of consensus surrounding the most suitable methods for GSA. The literature contains dozens of tools^10–18^ for performing gene set analysis, but there is little agreement on how to choose between so many competing ideas^19–21^. Existing GSA methods are frequently cited for flaws including insufficient power^22^, an inability to provide statistically valid tests under certain parameter settings^23^, and a reliance on permutation-based inference^24^. More specifically, many existing methods fail to control the Type I error rate for genes with unconventional characteristics - for example, genes with a small number of SNPs or a large amount of correlation^25,26^. Permutation offers a valid solution, but complicated resampling schemes often muddle the null hypotheses being tested and result in confusing interpretations^18,27^. Permutation can also be extremely computationally expensive when attempting to control for multiple testing, e.g. a Bonferroni correction to control the family wise error rate over 10,000 tested pathways requires approximately ten million iterations.

Another key challenge, which few methods have attempted to address, is the lack of standardized gene set lists in the public domain^28,29^. While there exist multiple sources for such information, gene set definitions can be highly differentiated across databases, and small differences can lead to large inconsistencies in results. As a representative example, we consider the case of the *FGFR2* gene in breast cancer. In the largest breast cancer GWAS cohort to date, SNPs near *FGFR2* demonstrate association at *p* < 10^−300^; these are the smallest p-values across the entire genome. Subsequent pathway analysis of this GWAS^30^ tests approximately 4,000 pathways - 182 containing *FGFR2* - and concludes that 86 of the top 100 pathways all contain *FGFR2*. Clearly a pathway is extremely likely to be found significant if it contains *FGFR2*, and thus pathways unrelated to breast cancer may be artificially driven to the top of the results based on the decision to include or exclude a single gene. For instance, the Gene Ontology^31^ pathway Ear Morphogenesis includes *FGFR2* and is ranked among the top 100 most significant gene sets, but as we will show later, the same pathway defined without *FGFR2* possesses minimal association with breast cancer. Pathway lists that do not include *FGFR2* in their version of an ear morphogenesis pathway may have difficulty replicating a seemingly strong association.

In this paper we make three key contributions toward overcoming the above challenges. First, we introduce a class of supremum-based goodness-of-fit tests for gene set analysis, demonstrating how they can be adapted for use in the GSA framework and focusing in particular on use of the Generalized Berk-Jones (GBJ) statistic. Originally developed for optimal detection of sparse signals in independent data^32^, the aforementioned class includes the Higher Criticism (HC) and Berk-Jones (BJ) statistics, which have been adapted for correlated data through the Generalized Higher Criticism (GHC) and GBJ. These statistics possess, in a certain sense, optimal power for detection of a set-based effect when many elements of the set may individually demonstrate no association, as in testing for the effect of a gene set that may contain an appreciable subset of neutral variants. Among tests derived from this class of statistics, GBJ demonstrates more robust and powerful performance than the Generalized Higher Criticism^33^ when testing gene sets in a range of practical settings^34^, while other tests in the class have not yet been adapted to account for correlation. Unlike other GSA methods that rely on permutation to adjust for correlation and frequently summarize information at the gene level with a single value^28^, GHC and GBJ admit analytic p-value calculations that avoid permutation and automatically account for features such as the size of a gene set and LD patterns between SNPs. Both tests have also been shown to protect Type I error rates at very stringent levels^34^.

Secondly, we propose a step-down GSA inference procedure that can identify gene sets driven to significance solely by a few genes, as opposed to gene sets containing signals spread throughout the entire group. This procedure relies on the self-contained^6^, one-step nature of GBJ, which allows for a unified approach to testing the association between single genes and the outcome. Re-analyzing a set after removal of its most significant genes can uncover the gene sets and themes which will still demonstrate replicable associations over different pathway databases. Such sets are also arguably more important from a biological standpoint.

Thirdly, we illustrate the utility of both GBJ and the step-down procedure through simulation and re-analysis of summary statistics from a large breast cancer GWAS dataset. Simulation demonstrates the additional power of GBJ over alternatives including GHC, the popular Gene Set Enrichment Analysis (GSEA*)^11^*, and MAGMA^18^. Simulations also show that the step-down procedure is adept at distinguishing between pathways with single-gene signals and those containing multi-gene signals. Review of the breast cancer step-down results illustrates that many seemingly significant sets are completely dependent upon *FGFR2* and a few other genes for their strong association signals. These sets deserve further screening before their significant association with breast cancer is reported.

Finally, we apply GBJ in a cross-phenotype analysis of breast cancer, height, and schizophrenia to investigate certain gene set properties that have received less attention in the literature. This work finds that the pathways significantly associated with human height are much more likely to contain signals spread throughout an entire gene set, while pathways associated with breast cancer are more likely to see their signals localized to a few genes. We additionally observe that immune system pathways are highly associated with schizophrenia, while growth pathways are often linked with height and breast cancer.

## Materials and Methods

### Overview of GBJ

The Generalized Berk-Jones statistic provides a powerful parametric approach for testing the association between a set of SNPs and a phenotype using marginal SNP summary statistics. Consider a gene set that contains *d* SNPs. The *d* summary statistics for these SNPs, **Z** = (*Z*_1_, …, *Z_d_*)^T^, follow a joint multivariate normal distribution,

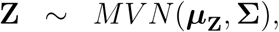

where the diagonal elements of **Σ** are all 1. GBJ aims to robustly test *H*_0_: ***μ*_Z_** = **0**_*d*×1_ against *H*_1_: ***μ*_Z_** ≠ **0**_*d*×1_ when ***μ*_Z_** contains a subset of zeros and while accounting for the correlation between test statistics. Thus common GSA features such as LD between SNPs and multiple neutral variants are automatically incorporated into the statistical framework. GHC and other tests in the goodness-of-fit class operate similarly, but for reasons of space, we will limit comparisons to simulation results.

The null hypothesis corresponds to the situation where no SNPs in the entire set are associated with the outcome, after correction for confounders. When performing GSA with genotype-level data for each subject, it is necessary to both calculate **Z** and estimate **Σ**, and when summary statistics are available, it is only necessary to estimate **Σ**.

### Calculation of **Z** and Σ with genotype-level data

Suppose we have genotype-level data for *i* = 1, 2, …, *n* subjects at a set of *j* = 1, 2, …, *d* SNPs, so that the genotype vector for subject *i* is **G**_*i*_ = (*G*_*i*1_, …, *G_id_*)^*T*^. Let **G** = [**G**_1_, …, **G**_*n*_]^*T*^ be the *n* × *d* genotype matrix. Suppose also that we have a set of *q* additional covariates contained in **X** = [**X**_1_, …, **X**_*n*_]^*T*^, which is an *n* × *q* matrix with **X**_*i*_ = (*X*_*i*1_, …, *X_iq_*)^*T*^ for *i* = 1, …, *n*. Denote the outcome by *Y_i_* and let *μ_i_* be the mean of *Y_i_* conditional on **G**_*i*_ and **X**_*i*_. Consider the generalized linear model^35^ (GLM) for *μ_i_* given by

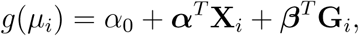

where *g*(⋅) is a canonical link function, for instance, *g*(*μ_i_*) = *μ_i_* for normally distributed phenotypes and *g*(*μ_i_*) = logit(*μ_i_*) for binary phenotypes. We are interested in testing the null hypothesis of no gene set effect *H*_0_: ***β*** = **0**_*d*×1_ against the alternative *H*_1_: ***β*** ≠ **0**_*d*×1_. As some SNPs in a gene set are likely neutral variants, we allow elements of ***β*** to equal 0 under *H*_1_. The marginal score test statistic under the null is

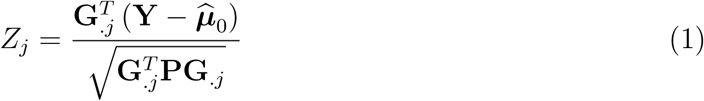

for any SNP *j* = 1, …, *d* in the gene set. Note that *H*_0_: ***β*** = **0**_*d*×1_ in the regression model corresponds to the null hypothesis *H*_0_: ***μ*_Z_** = **0**_*d*×1_ from above. Here **Y** = (*Y*_1_, .., *Y_n_*)^*T*^, ***μ̂***_0_ = 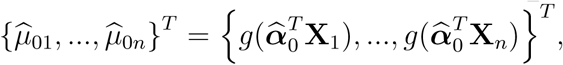 and ***α̂***_0_ is the MLE of ***α*** under *H*_0_: ***β*** = **0**_*d*×1_. Also we define the single variant vector **G**_.*j*_ = (*G*_1*j*_, …, *G_nj_*)^*T*^, the projection matrix **P** = **W**– **WX**(**X**^*T*^ **WX**)^−1^**X**^*T*^ **W**, and the GLM weight matrix **W** = diag {*a*_1_(*ϕ̂*)*v*(*μ̂*_01_), …, *a_n_*(*ϕ̂*)*v*(*μ̂*_0*n*_)}. The standard GLM dispersion parameter is given by *a_i_* (*ϕ̂*), and the standard GLM variance function is *v*(*μ̂*_0*i*_), where *v*(*μ̂*_0*i*_) = 1 for a normally distributed phenotype and *v*(*μ̂*_0*i*_) = *μ̂*_0*i*_(1 − *μ̂*_0*i*_) for a binary phenotype. The *Z_j_* are asymptotically equivalent to the individual SNP test statistics calculated by many popular tools, such as the Wald statistics produced by PLINK^36^. A consistent estimate of the correlation between *Z_j_* and *Z_k_* is then given by

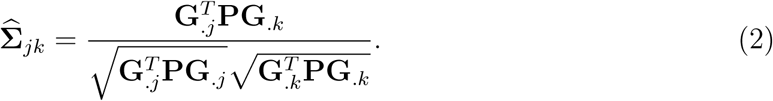

### Estimation of Σ with precalculated summary statistics

Precalculated marginal summary statistics are much more available than subject-level data. When using these summary statistics, we need to estimate their correlation structure from a reference LD panel containing individuals of a similar ethnicity, e.g. data from the 1000 Genomes Project^37^. Specifically, let 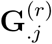 and 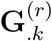 denote the genotype of each subject in the reference panel at SNPs *j* and *k*. Also let **X**^(*r*)^ = (**1**, **PC**_1_, …, **PC**_*m*_) denote a modified design matrix, where *m* is the same number of principal components used in the original analysis, and **PC**_1_,.., **PC**_*m*_ are PCs calculated from the reference data. Using **X**^(*r*)^ instead of 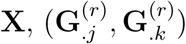 instead of (**G**_.*j*_, **G**_.*k*_), and any constant in place of 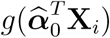 in Equation 2 provides a good approximation to **Σ̂**_*jk*_.

The motivation for this approximation comes from the observation that the primary confounders of the SNP-outcome relationship are the principal components. Thus a close substitute for the original design matrix in Equation 2 can be constructed using only the PCs. In practice we have found this substitution to be very reliable.

### The Generalized Berk-Jones statistic

Next assume we have calculated or been supplied a vector of test statistics 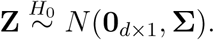 Denote by Φ̄(*t*) = 1 − Φ(*t*) the survival function of a standard normal random variable, with Φ^−1^(*t*) denoting its inverse. Further designate ∣*Z*∣_(*j*)_ as the order statistics of the vector that arises from applying the absolute value operator to each element of **Z**, so that ∣*Z*∣_(1)_ is the smallest element of **Z** in absolute value. Finally define the significance thresholding function

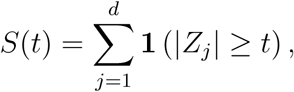

where *t* > 0 is the threshold.

It is helpful to think of *S*(*t*) as the “number of significant SNPs at threshold *t*.” For example, in GWAS, some researchers set *t* = 5.45131 so that *S*(*t*) counts the number of SNPs with p-values less than 5 × 10^−8^, the commonly-used cutoff for declaring genome-wide significance^38^. Other set-based methods implicitly set *t* equal to ∣*Z*∣_(*d*)_ and carry forward ∣*Z*∣_(*d*)_ as the representative test statistic for the entire set^23^. However in both of these examples, the choice of *t* is rather arbitrary and relies on a one-size-fits-all-sets approach. In particular, neither choice of *t* makes full use of the data, ignoring factors such as the size of the set and the LD pattern among SNPs. More importantly, moderately significant SNPs that do not reach a GWAS threshold or demonstrate the lowest p-value in a gene can cumulatively produce a major contribution to the phenotype. A key concept behind GBJ is that it can adaptively find the threshold *t* that best maximizes power for any given set while adjusting for the size of the set and the correlation structure of the SNPs.

Consider first the case of no **LD**, **Σ** = **I**, where **I** is the identity matrix. Then for a fixed *t* under the null hypothesis, *S*(*t*) has the binomial distribution *S*(*t*) ~ *Bin*(*d*, *Π*) with *Π* = 2**Φ̄**(*t*). This observation motivates the Berk-Jones (BJ) statistic^39^, which can be written as^40^,

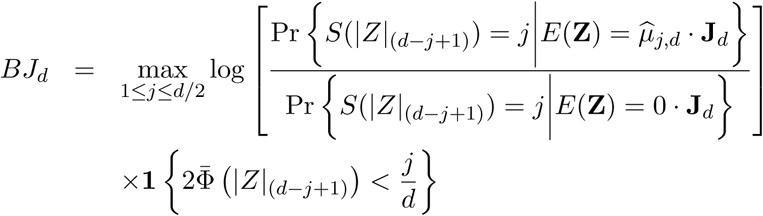

where 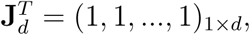 and *μ̂*_*j*,*d*_ > 0 solves the equation

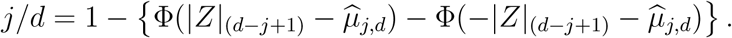

In other words, Berk-Jones is the maximum of a set of likelihood-ratio tests performed on *S*(*t*) at all observed test statistic magnitudes greater than or equal to the median observed magnitude. By taking the maximum over these different thresholds, Berk-Jones allows the data to set the threshold which provides the most power in the presence of an appreciable subset of neutral variants.

When SNPs in a gene set are in LD and **Σ** ≠ **I**, then *S*(*t*) no longer has a binomial distribution, and the Berk-Jones statistic can lose much of its power in finite samples. However the Generalized Berk-Jones incorporates the additional correlation information by explicitly conditioning on **Σ**,

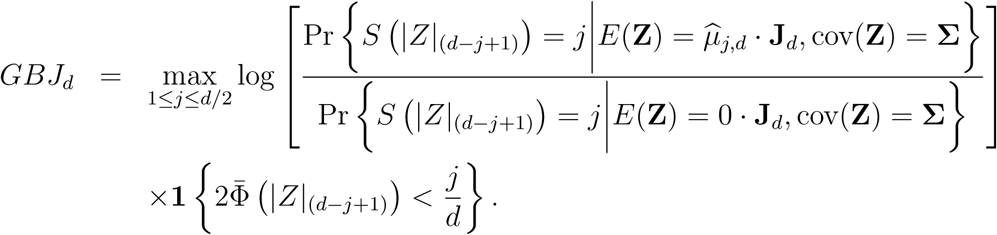

GBJ is still the maximum of a set of likelihood ratio type tests, but it gains notable power over BJ by accounting for the correlation between test statistics. When **Σ** = **I**, GBJ reduces to the standard Berk-Jones. The p-value of the GBJ statistic can be calculated analytically.

In integrating the test statistic for each SNP in the gene set, GBJ incorporates much more information than methods which only keep the most significant p-value in each gene^11,14,16^ and discard the rest in an ad-hoc fashion. Other tests may not discard individual test statistics but instead summarize the values in mean^12^ or count-based procedures^14,15^ so that the individual magnitudes are lost. For example, test statistics of *Z*_1_ = −2.5 and *Z*_2_ = 5 make equal contributions to a procedure that counts the number of p-values less than 0.05. However, the variant with test statistic *Z*_2_ = 5 clearly conveys more information about genotype-phenotype association than the variant with *Z*_1_ = −2.5. In contrast, GBJ does not discard or summarize information and utilizes each marginal test statistic, as well as the joint correlation structure. When **Σ** = **I**, GBJ enjoys certain asymptotic power properties that other tests may not demonstrate^40^.

### GBJ step-down inference

As an extension of GBJ, we propose a step-down inference procedure to filter out gene sets that are driven to significance based on the signal from only a very small proportion of genes, such as the earlier example involving Ear Morphogenesis and *FGFR2*. The procedure begins by performing gene-level association analysis. First, create a list of the unique genes over all gene sets under consideration, then define each gene as its own set and apply GBJ over each single gene to find single gene p-values. Sort the single genes in increasing order of p-value, which can be interpreted as ranking the genes by their level of association with the outcome.

For any given gene set, obtain a measure of how much its association signal is dependent on a few highly associated genes by applying GBJ to the set after removing all SNPs that belong to the top *k* genes. Setting *k* = 1 will identify gene sets where the signal is driven by a single gene. If a gene set remains significant even after removing *k* > 1 genes, then the set possesses signals dispersed through many different genes. In this work we will use *k* = 1 and *k* = 3. As we show below, *FGFR2* drives the significance of many gene sets in a breast cancer analysis, but the step-down procedure allows us to uncover pathways that show no association outside of *FGFR2* and may be less suitable for follow-up.

### GWAS summary statistics datasets

Three large, publicly available summary statistic datasets are analyzed in this study. We first obtained summary statistics from the largest breast cancer GWAS cohort to date^30^, with 122,977 cases and 105,974 controls of European ancestry. Most subjects were genotyped on the OncoArray, a custom-designed array for cancer studies that also has genome-wide coverage of over 570,000 SNPs. Data for these subjects were imputed using the full 1000 Genomes Project Phase 3 reference panel, resulting in estimated genotypes for approximately 21 million variants. Other subjects were included from various smaller studies, including the iCOGS project^41^ and 11 smaller GWAS. Results across studies were then meta-analyzed, and after quality control, approximately 12 million SNPs produced a final test statistic for association with breast cancer.

For height, we downloaded summary statistics from the Genetic Investigation of Anthropometric Traits (GIANT) GWAS^42^. In this study, 253,288 individuals of European ancestry were genotyped on multiple Affymetrix, Illumina, and Perlegen arrays. All individuals were then imputed to the Phase II CEU HapMap release. After meta-analysis and quality control, there were over 2.5 million SNPs with a final summary statistic for association with height.

The last dataset used was downloaded from the Psychiatric Genomics Consortium schizophrenia mega-analysis^43^. In the primary GWAS of this study, 34,241 cases and 45,604 controls were genotyped across 49 cohorts. The vast majority of samples were obtained from subjects of European descent, but three cohorts did contain individuals of East Asian ancestry. All subjects were imputed using 1000 Genomes Project data as a reference panel, and test statistics were meta-analyzed across cohorts. For this study, summary statistics were made available for approximately 9.5 million variants.

### Pathway definition database

To provide a fair comparison between GBJ and GSEA in re-analysis of the breast cancer data, we conduct all pathway analysis using the same gene set database (Gary Bader Lab, Human GOBP all pathways, no GO IEA; April 1, 2017) used in the original breast cancer pathway analysis^30^. This file compiles gene sets from a number of databases including Gene Ontology (GO) Biological Process^31^, Reactome^44^, Panther^45^, and others. In all, the file contains 16,528 gene sets. While we are unaware of a comprehensive method to assess the quality of gene set databases, advantages of this list include the incorporation of multiple different sources, public availability, and monthly updates.

As the selected pathway database is a direct aggregation of multiple sources possessing varying levels of curation, some preprocessing of the list is necessary before beginning analysis. We first truncate the database to remove all pathways with more than 200 genes or less than three genes. Removing pathways with a large number of genes is common practice^46,47^, as extremely large pathways are difficult to interpret, and this step was performed in the original GSEA analysis as well. The original GSEA analysis further removed all pathways with fewer than ten genes, which is also a relatively common strategy to reduce false positives and lower the multiple testing burden^48^, however it has been noted that this threshold may exclude certain specific and informative functional sets such as protein complexes^26^. We choose to set the lower limit for pathways at three genes because there may be interesting insights to be gleaned from smaller gene sets and because we believe GBJ is powerful enough to overcome the increased multiplicity burden. In total, there are 10,742 pathways with between three and 200 genes.

### SNP-gene mapping

For each set of summary statistics, we map individual SNP test statistics to gene sets if they lie within 5 kb of a gene in the set, with coordinates provided by Ensembl 90 gene annotations^49^. Estimates of the correlation between summary statistics are calculated using unrelated subjects from European cohorts (TSI, FIN, GBR, IBS, CEU) in the 1000 Genomes Phase 3 data release. Summary statistics belonging to SNPs that have minor allele frequency less than 3% in the reference panel are removed as their data would be unstable for estimation. The original GSEA analysis maps SNPs to genes in a slightly different manner and also performs some additional manual curation. These additional steps are unique to the breast cancer dataset, so to preserve generality and facilitate comparison of results across traits, we do not include them here.

To further reduce the computational burden of a GBJ analysis, we additionally trim SNPs that are in the same gene and are correlated at *r*^2^ > 0.5. This pruning is performed in PLINK and occurs at successive multiples of 0.5 for very large sets, so that all tested sets are less than 1,500 SNPs in size. Even after the pruning procedure, GBJ still incorporates a very large amount of information, as the median number of SNPs in our breast cancer analysis is 470. In contrast, the median number of genes in the GSEA analysis is 26; a test on 26 genes with GSEA incorporates only the 26 minimum p-values from those genes.

The data-intensive nature of GBJ does pose problems for a small number of extremely large pathways. Some gene sets are larger than 1,500 SNPs even after pruning all pairs of SNPs in the same gene with *r*^2^ > 0.0625. These sets are not tested to remain consistent in our analysis protocol across all three phenotypes. However for typical GSA focusing on a single outcome, it would be straightforward to perform additional pruning or manual curation to accommodate testing the largest pathways.

### Simulation

We use simulation to compare the performance of GBJ against the self-contained version of GSEA (as described by Wang et al.^11^), self-contained MAGMA, and GHC. GSEA and MAGMA were two of the best-performing methods in the comprehensive simulation study of a recent GSA review^28^. To match the settings of our real data analysis, we use real genotype data from the reference panel of *n* = 350 unrelated Europeans in the 1000 Genomes Project. These genotypes have been pruned as described above, and we use all SNPs located in 10,000 genes chosen at random. Each of the 10,000 genes contains between 7 and 25 SNPs, which corresponds to the middle 50 percentile of pruned gene size over the entire gene database.

For each iteration of the power simulation, we choose 10 genes at random to be the tested pathway, and *a* of the 10 genes are given *b* causal SNPs each. The true disease model is then

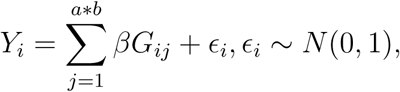

where *G_ij_* is the genotype of subject *i* at causal SNP *j*. Marginal summary statistics for each SNP in the pathway are calculated according to Equation 1 and their covariance is estimated according to Equation 2. We study the four conditions (*a* = 8, *b* = 1), (*a* = 4, *b* = 2), (*a* = 2, *b* = 4), and (*a* = 1, *b* = 4), and we vary the causal SNP effect size *β* from 0 to 0.3. When *β* = 0 we are estimating the Type I error rates of the tests, and when *β* > 0 we are comparing their power. MAGMA is applied with default self-contained settings, and 1000 permutations are used for GSEA inference. We test at *α* = 0.05 with 100 runs performed at each parameter setting.

We also perform a simulation to assess the validity of the step-down inference procedure. Data is generated as in the power simulation, but in each iteration we remove the most significant gene - chosen separately for each test - from the pathway before generating a pathway p-value. For GBJ, GHC, and MAGMA we use the default gene-level analysis to determine the significance of each gene, and for GSEA we remove the gene with the most significant SNP. In the simulation with (*a* = 1,*b* = 4) we are benchmarking the discriminatory ability of this procedure, as the step-down procedure should remove the only causal SNPs, resulting in power equal to the Type I error rate regardless of effect size. In settings with *a* > 1, we can assess power to identify gene sets with dispersed signals.

### Classification of biologically important systems

To summarize our results from applying GBJ across multiple phenotypes, we search for the biological systems that demonstrate the largest degrees of significance across each phenotype. Pathways are categorized into different systems by exploiting the directed acyclic graph structure of Gene Ontology (GO) Biological Process pathways. Starting with the top-level Biological Process category, GO defines successively smaller groups of pathways so that each child term is more specialized than its parent term. Using this natural structure, it is possible to group categories of pathways at different levels of granularity.

We create categories from the first level immediately following the Biological Process root. Specifically, we use the 11 top-level sets Biological Adhesion, Cellular Component Organization or Biogenesis, Developmental Process, Growth, Immune System Process, Localization, Locomotion, Metabolic Process, Reproduction, Response to Stimulus, and Signaling. For a given phenotype and category, we first calculate the expected number of significant pathways in the category conditional on the total number of significant pathways for the outcome. If pathways in each category truly have the same chance of reaching significance, then the expectation is simply equivalent to the percentage of all tested pathways that belong to the category multiplied by the total number of pathways significantly associated with the outcome. For each phenotype, we calculate the difference between the observed and expected number of significant pathways arising from each category, taken as a percentage of the expected number, to determine which categories harbor more significant pathways than expected. Pathways are deemed significant using the Bonferroni-corrected significance level of *α* = 0.05/10, 742 = 4.65 ⋅ 10^−6^.

## Results

### Simulation

We first compared the power of GBJ to other gene set methods through simulations carried out with genotypes from the 1000 Genomes Project. Because non-GSEA tests utilize data from multiple SNPs in each gene, we expected such tests to perform better at detecting pathways with many medium-sized signals that GSEA must discard. However, even when each gene held only one signal, a situation that would appear very favorable for GSEA, we found that the power of GBJ was either the largest or approximately the largest at each effect size (Fig 1A). When signals were more densely packed into a smaller number of genes (Fig 1B-D), GBJ and GHC increased their advantage over GSEA and MAGMA significantly, demonstrating the power of the goodness-of-fit approaches over a variety of sparsity settings. Self-contained GSEA was the clear third-best performer, generally demonstrating more power than self-contained MAGMA. We also observed in this simulation that power for GBJ and GHC improved as more signals were placed in the same gene (as opposed to spread throughout different genes), because increased correlation between signals can increase the power of these tests^34^.

**Figure 1:**
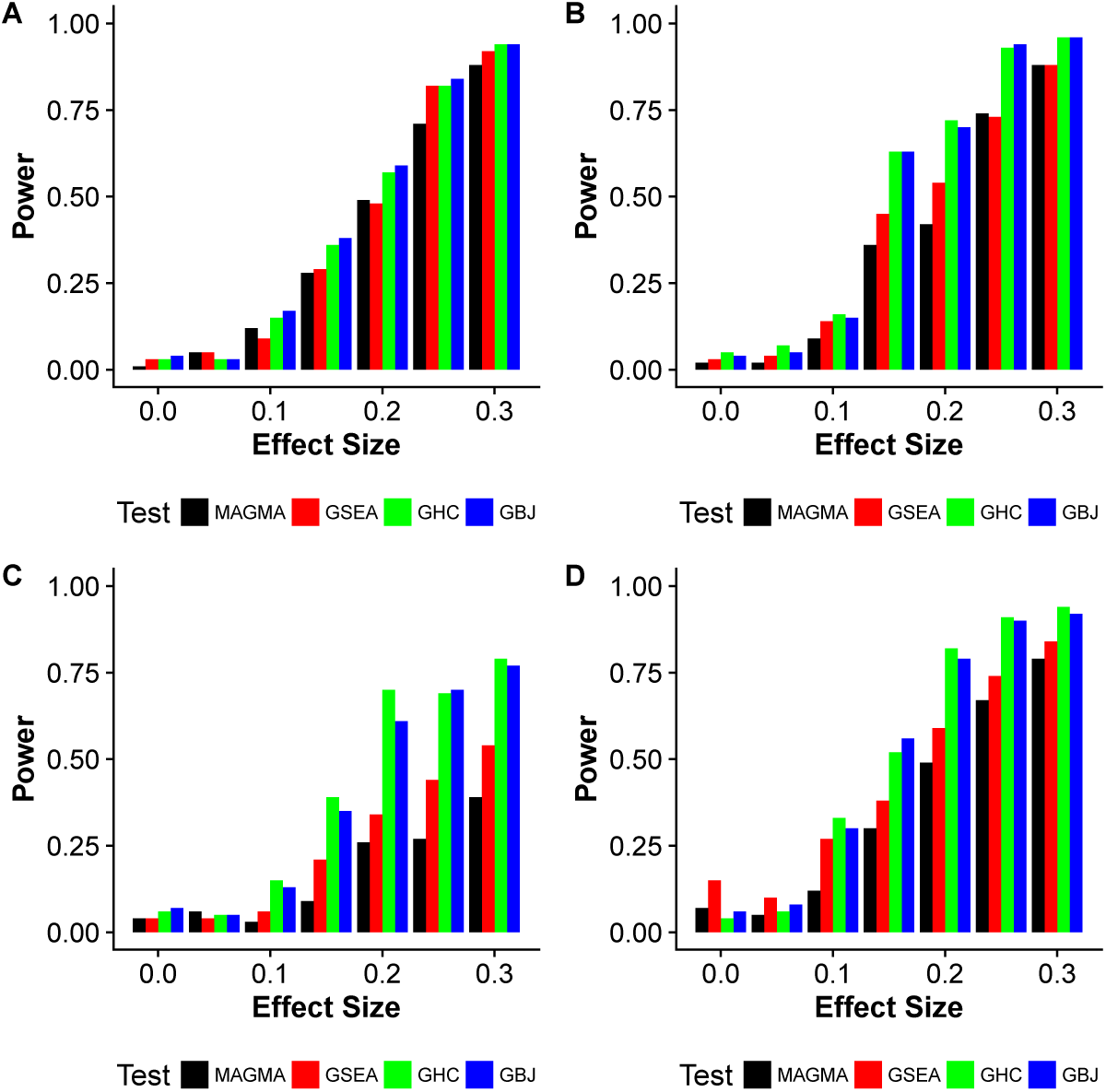
GSA power simulation over four different configurations of gene signal density. Simulated power of MAGMA, GSEA, GHC, and GBJ (all self-contained versions) with random sets of ten genes selected from 10,000 total genes. From the ten genes in the set, *a* genes are selected to hold *b* causal SNPs each. The four subfigures correspond to (A) *a* = 8, *b* = 1, (B) *a* = 4, *b* = 2, (C) *a* = 1,*b* = 4, (D) *a* = 2, *b* = 4. The effect size is given on the x-axis. We perform 100 simulations at each parameter setting and test at *α* = 0.05. GBJ and GHC increase their power advantage over GSEA and MAGMA as the same number of signals are located in fewer genes.

Additional simulations showed similar trends for a different number of background genes (S1 Fig). Power for GHC and GBJ was often close, but in general, we would expect GBJ to outperform GHC in GSA, as set sizes in GSA are relatively large and are likely to contain a moderate number of signals. GHC would be a better choice under extreme sparsity, as in only one or a few signals in the entire set.

Both GHC and GBJ appeared to control the Type I error rate fairly well at *α* = 0.05, as their power remained approximately equal to the nominal size of the test when there was no true effect. This observation suggested that our inference was valid for the situations considered in simulation. Comparable results also held for a different number of background genes (S3 Fig).

Simulations for the step-down procedure (Fig 2) showed that the proposed method was indeed able to recognize pathways containing signal in only one gene, regardless of the choice of GSA test statistic. When only one gene contained causal SNPs (Fig 2C), the power for all tests remained around the Type I error rate regardless of effect size, as the causal SNPs were removed before pathway-level inference. When multiple genes contained causal SNPs, GBJ outperformed the other tests again in a majority of the situations, with GHC a close second. While the step-down procedure showed slight power loss compared to the standard analysis of Fig 1, the difference was not dramatic, and the segregation of single-gene effects was very good. Thus we suggest the step-down procedure as an important complementary tool in GSA.

**Figure 2:**
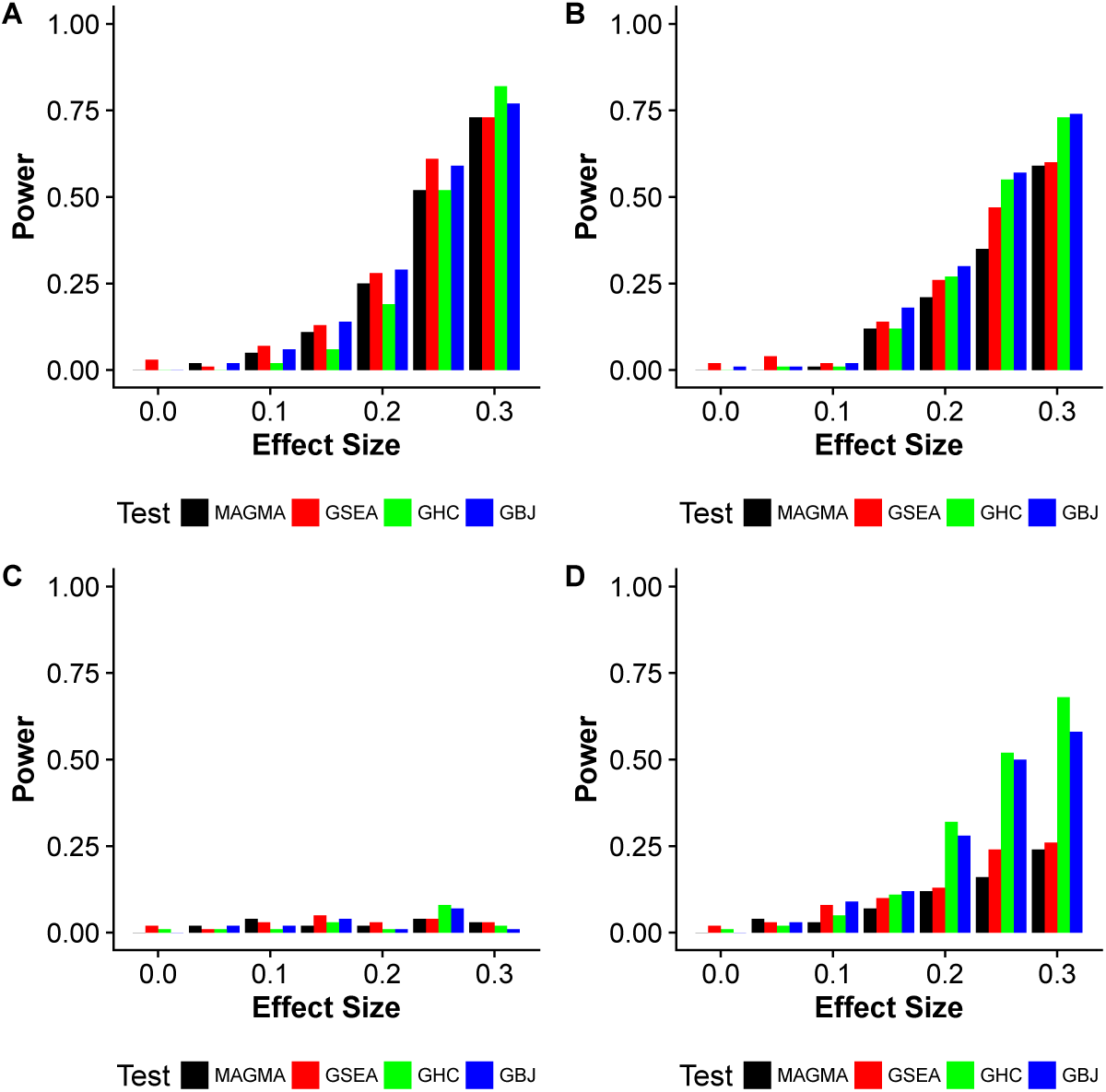
Step-down inference power simulation over four different configurations of gene signal density. Simulated power using step-down inference procedure with MAGMA, GSEA, GHC, and GBJ (all self-contained versions) with random sets of ten genes selected from 10,000 total genes. From the ten genes in the set, *a* genes are selected to hold *b* causal SNPs each. The four subfigures correspond to (A) *a* = 8, *b* = 1, (B) *a* = 4, *b* = 2, (C) *a* = 1, *b* = 4, (D) *a* = 2, *b* = 4. The effect size is given on the x-axis. For each method and each iteration, we first determine a most significant gene. Then that gene is removed and we perform inference on the remaining nine genes in the set. We perform 100 simulations at each parameter setting and test at *α* = 0.05.

### Re-analysis of breast cancer GWAS

We next investigated whether the same trends seen in our simulation could be seen in re-analysis of the breast cancer summary statistic dataset. Self-contained GSEA was originally used to analyze a total of 4,507 pathways containing more than ten genes, finding 448 to be significant when using permutation to control the false discovery rate (FDR) at *q* = 0.05. GBJ was applied to 10,742 pathways (with more than three genes) from the same master list and found 2,703 to be significant at the Bonferroni-corrected family wise error rate of 4.65⋅10^−6^. When we restricted comparisons to the 3,952 pathways tested by both approaches, GSEA found 352 significant while GBJ found 2,095 significant at their respective error rates.

From the raw significance numbers alone, GBJ appeared to offer far more power in a real GWAS summary statistic dataset. GBJ found more than twice as many significant pathways as GSEA on a percentage basis, even when controlling a more stringent error rate and using a conservative Bonferroni correction. The increased power could not be attributed to smaller pathways alone, as GBJ declared a higher percentage of pathways significant across various pathway sizes (S2 Fig). To further investigate, we compared the GSEA and GBJ significance ranking (Fig 3) of the 3,952 pathways tested by both methods (p-value or q-value rank out of 3,952, lower is more significant). To emphasize the role of moderately strong associations, points were colored according to their density of suggestive signals, which we defined as a pathway’s proportion of SNPs with *p* < 10^−5^. SNPs demonstrating such a level of association would generally not stand out as the strongest signal in their region, but a large proportion of suggestive signals in any single gene or pathway could still indicate biologically relevant gene sets. For the sake of presentation, we only plotted pathways ranked in the top ten percentile by GSEA, in the top ten percentile by GBJ, in the 30^*th*^-40^*th*^ percentile by GBJ, and in the 60^*th*^-70^*th*^ percentile by GBJ (see S3 Fig for full data).

**Figure 3:**
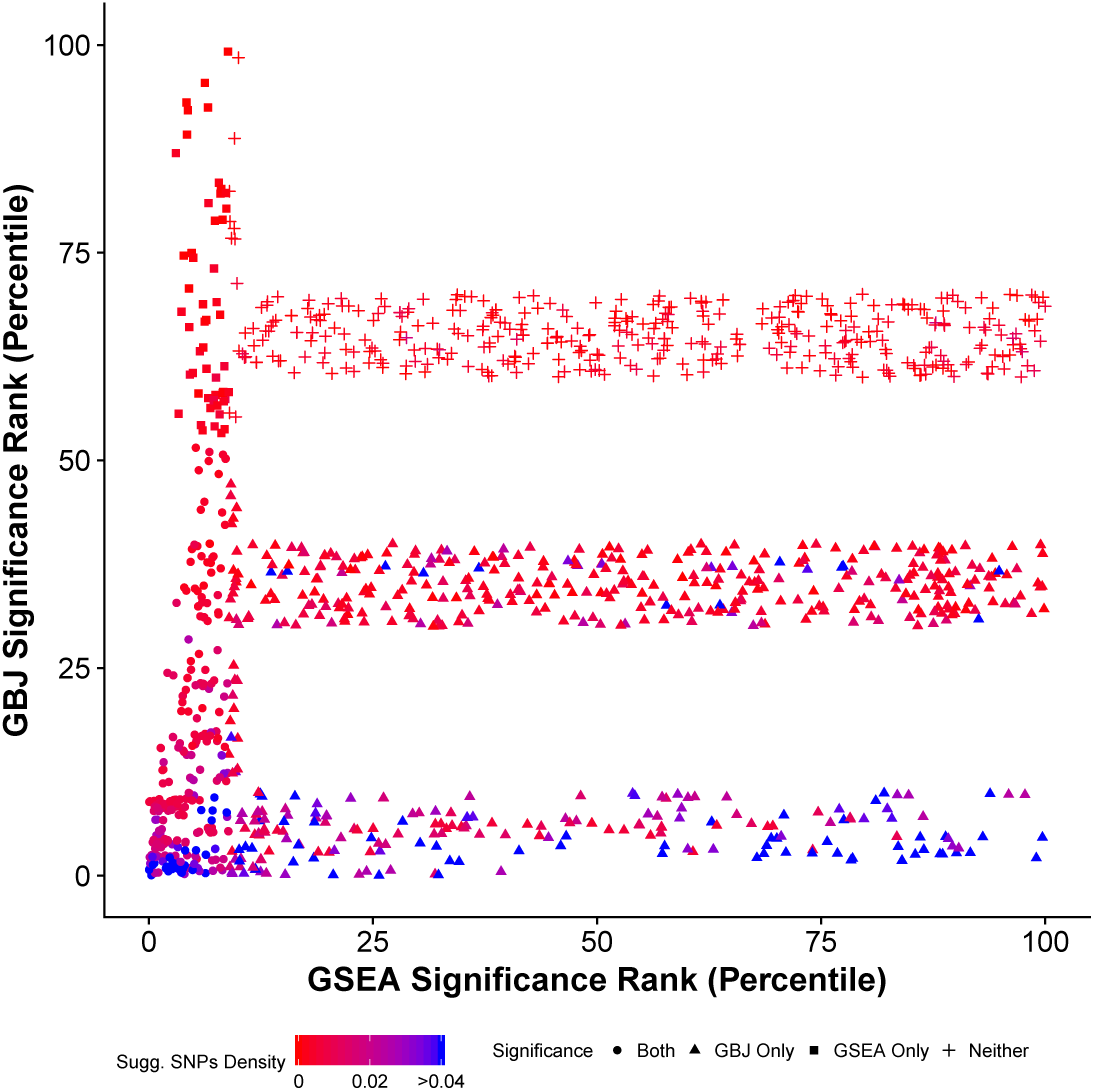
Significance rank of breast cancer pathways tested by both GBJ and GSEA. The significance ranking of pathways according to GBJ and GSEA, colored according to density of SNPs with *p* < 10^−5^. Pathways are ordered according to p-value (GBJ) or q-value (GSEA), smaller rankings indicate more significance. For the sake of presentation, we only show pathways in the top ten percentiles of either GSEA or GBJ as well as pathways in the 30^*th*^-40^*th*^ and 60^*th*^-70^*th*^ percentiles of GBJ. Pathways ranked by GBJ as more significant generally have a higher proportion of SNPs with *p* < 10^−5^. In contrast, there is no discernible relationship between a pathway’s GSEA ranking and its density of suggestive signals.

The frequency of blue pathways - indicating higher density of suggestive signals - clearly increased for pathways that were ranked as very significant according to GBJ. Such a pattern was desirable, as pathways with a higher density of small p-values should be more significant. However, scanning horizontally across the plot, there did not appear to be a strong relationship between GSEA rank and frequency of blue pathways. Approximately the same number of blue pathways could be found near the GSEA 25^*th*^, 50^*th*^, and 75^*th*^ percentiles. This pattern suggested that GSEA was not very sensitive to the density of medium-strength signals.

The proportion of SNPs with *p* < 1 ⋅ 10^−5^ is not a perfect measure for evaluating the significance of pathways, as other factors including linkage disequilibrium and strength of the largest signal do also play an important role. However, while GBJ takes into account all of the above factors, GSEA ignores signal density and disregards all p-values that are not the smallest in a gene. This difference was likely a major contributing factor to the large discrepancy between GSEA and GBJ rankings for points in the top left and bottom right hand corners of Fig 3. Other methods employing the same strategy of choosing a minimum p-value to represent each gene, for example, non-default versions of MAGMA, may possibly experience similar drawbacks.

### Single gene vs. set-based effects

Application of Generalized Berk-Jones to height and schizophrenia resulted in even more significant pathway findings than the breast cancer analysis (S1 Table). To uncover the gene sets that were driven to significance by only one or a few genes, we applied the GBJ step-down inference procedure (see Materials and Methods) for the top 500 pathways in each phenotype, all of which possessed an initial p-value of *p* < 1 ⋅ 10^−12^ (Fig 4). In a typical GSA, these pathways would likely receive the most attention for their extremely high levels of association.

**Figure 4:**
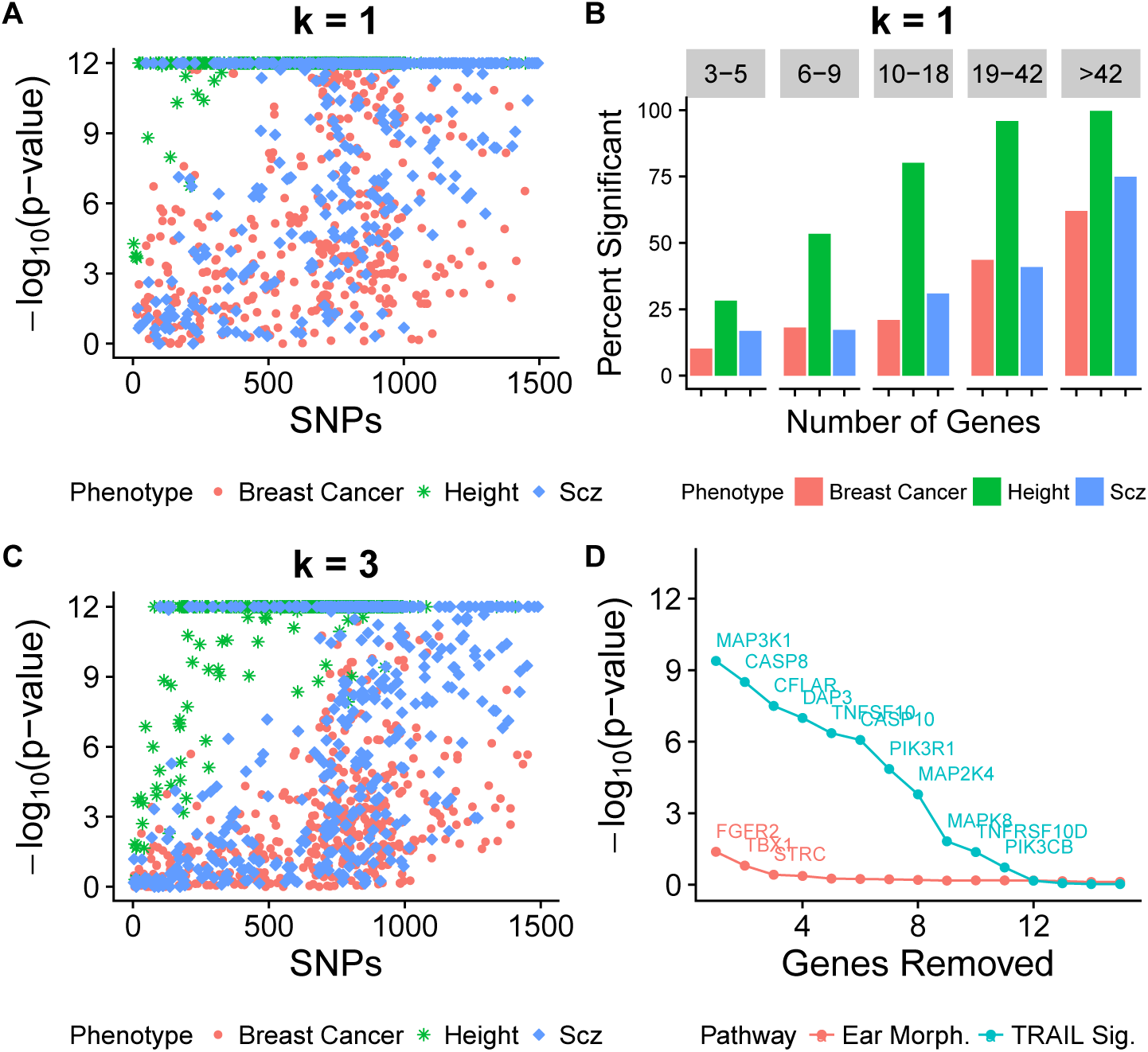
P-values of initially top-ranked pathways after removal of significant genes. (A) P-value after removal of most significant gene for top 500 pathways across each of three phenotypes. (B) Percentage of pathways with original association *p* < 1 ⋅ 10^−12^ passing Bonferroni-corrected significance threshold when top significant gene is removed, stratified by quantile of gene set size. (C) P-value after removal of three most significant genes for top 500 pathways across each of three phenotypes. (D) P-value of Gene Ontology Ear Morphogenesis and Nature Pathway Interaction Database TRAIL Signaling pathways as most significant genes are removed one by one (for association with breast cancer only). P-values less than 10^−12^ are truncated at this value. It appears that many of the most significant height pathways are driven to significance by multiple highly associated genes, while the opposite is true for breast cancer.

A large cluster of pathways from both breast cancer and schizophrenia dropped below the Bonferroni-corrected significance level after their most significant gene was removed, but breast cancer pathways appeared more highly concentrated at the bottom of the y-axis (Fig 4A), although some remained highly significant (S2 and S3 Tables and S4 Fig). Thus a single gene boosted evidence of association by many orders of magnitude for a large number of breast cancer pathways. In contrast, pathways associated with height generally remained significant even after removal of their most highly associated gene (Fig 4B). This trend persisted when removing the top three most highly associated genes from each pathway (Fig 4C). Only about 12% of breast cancer pathways survived the Bonferroni-corrected significance level after three genes were removed, while approximately 38% of schizophrenia pathways and 69% of height pathways still passed this threshold. A possible interpretation of these results could be that height was much more driven by pathway-level effects of many genes working together, while breast cancer risk factors were more localized to a few key genes. It is also possible that breast cancer signal was attenuated because the summary statistics included patients with multiple different subtypes, so signal may have been diluted compared to an ER-positive only or ER-negative only analysis.

Ear Morphogenesis was one example of a breast cancer pathway where the signal was almost entirely confined to a single gene, with an initial GBJ ranking of 100th most significant gene set (*p* < 1 ⋅ 10^−12^) before the step-down procedure. When *FGFR2* was removed from this pathway, the p-value of the modified gene set increased drastically to *p* = 0.041, far from the corrected significance level. As expected, the other genes were not very relevant to breast cancer; it is not recommended to further pursue replication of the set’s association, despite initially promising results from GBJ and GSEA. On the other hand, a gene set such as the Nature Pathway Interaction Database TRAIL Signaling pathway demonstrated more robustness to removal of its top gene, *MAP3K1*. TRAIL Signaling retained some set-level signal even as more and more top genes were removed from the gene set (Fig 4D). Along with *MAP3K1*, the pathway contained the significant genes *CASP8*, *CFLAR*, *DAP3*, and *TNFSF10*. All of these genes possessed single gene p-values less than 1 ⋅ 10^−5^ for association with breast cancer, while Ear Morphogenesis contained no such genes other than *FGFR2*. TRAIL Signaling as a mechanism has been studied extensively for its role in breast cancer^50^, supporting our finding of a pathway-wide effect that extends past the most significant genes.

Over the entire pathway database, 172 pathways containing *FGFR2* were tested for association with breast cancer, and 172 were significant at the Bonferroni-corrected threshold according to GBJ. Additionally, *FGFR2* was the most significant gene in 169 of these pathways. After removal of *FGFR2* from the pathway, only 71 of the 169 were still significant at the same threshold (S4 Table). Clearly, the composition of significant gene sets in any breast cancer pathway analysis will depend on the number of times *FGFR2* and select other genes (S5 and S6 Tables) appear in the pathway definition database.

### Significant biological systems

To summarize the results of our GBJ-based pathway analysis across three different phenotypes, we identified the biological processes where significant pathways in breast cancer, schizophrenia, and height were most likely to congregate (see Materials and Methods). In two reassuring results, we found that the percentages of significant pathways from the Growth category were higher than expected in breast cancer and height (Fig 5). Growth mechanisms have previously been found to play important roles in studies of breast cancer^30^ and height^3^. Another theme that has often been corroborated in the literature is the importance of the immune system in schizophrenia^43^. Immune-related pathways have been studied in connection with many psychiatric diseases, and our analysis underscored the reasons for such an approach, as we found a high density of significant schizophrenia gene sets arising from immune processes. Seeing that GBJ could identify outcome-category pairs known to be associated with each other offered further validation that our approach was selecting relevant pathways.

**Figure 5:**
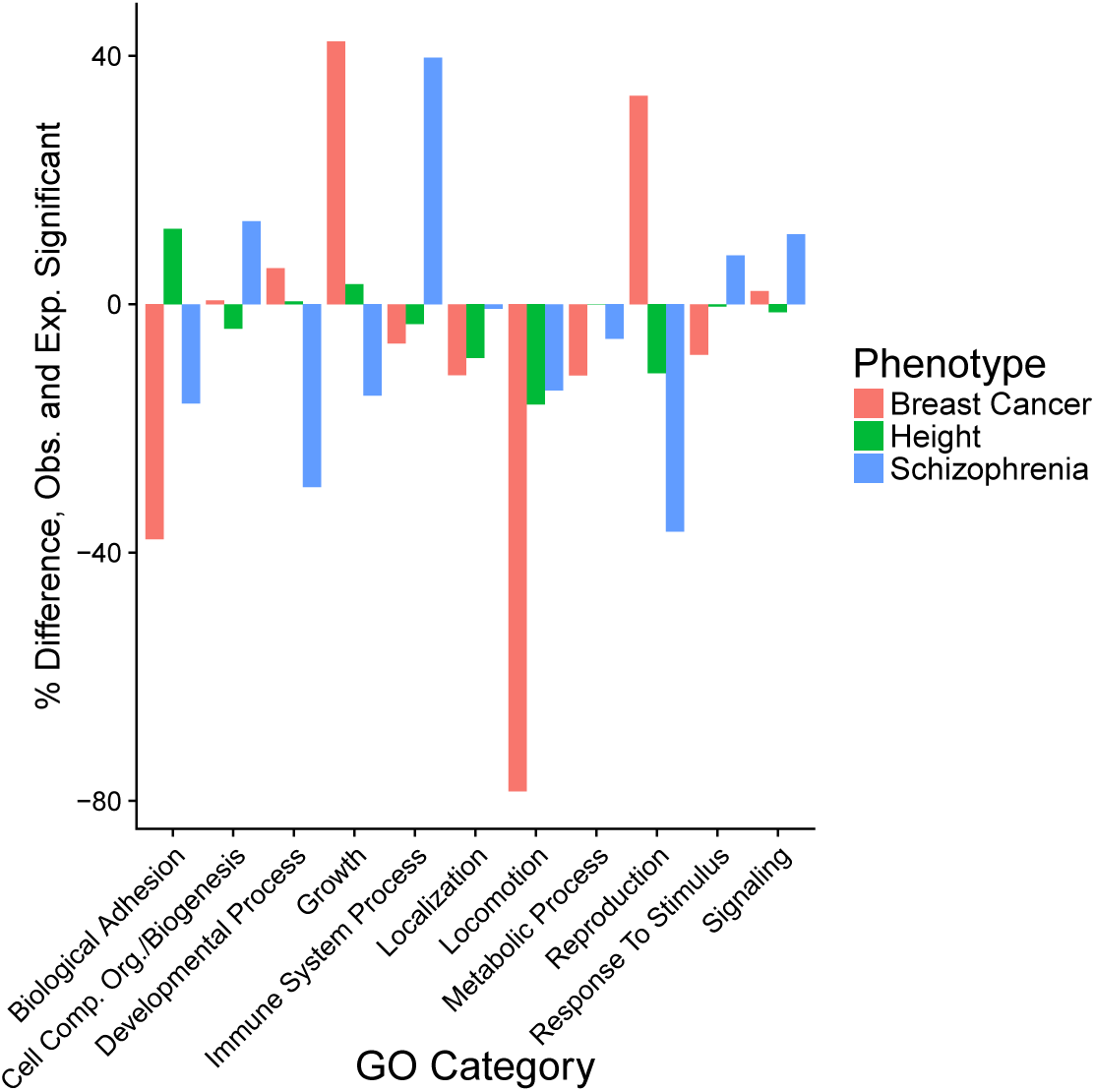
Difference between observed and expected number of significant pathways arising from Gene Ontology Biological Process categories. For a given phenotype, expected number is equal to the percentage of all tested pathways belonging to a category multiplied by the total number of significant pathways. The difference between observed and expected counts is expressed as a percentage of the expected number. A value greater than 0 indicates there are more significant pathways in a category than expected. A value less than 0 indicates there are less significant pathways than expected. A number of familiar themes are present, including the high number of significant height pathways related to growth and the high number of significant schizophrenia pathways related to the immune system.

On the other hand, GBJ also illuminated some outcome-category relationships that were not as widely familiar. For instance, we saw that there was a dearth of significant pathways related to schizophrenia in the Reproduction category. Thus there may be less benefit to searching for common drivers of risk between schizophrenia and breast cancer, which showed many more significant Reproduction pathways than expected. Similarly, all three phenotypes showed fewer than expected significant pathways in Locomotion, indicating that it may be more useful to prioritize other types of pathways when studying these outcomes. While negative findings are reported less often than their positive counterparts, these results still have the potential to inform researchers of the mechanisms that may not generate as many fruitful results.

## Discussion

Interest in GSA will likely continue to grow as more and more genotyping data is collected^28^, especially since single SNPs are still unable to explain much of the heritability in various phenotypes^51^. However, without appropriate statistical models to test for set-based effects, it will be difficult to correctly identify the gene sets that are truly associated with various outcomes. Many current GSA methods possess unknown operating characteristics and are difficult to interpret^18,29^. Our work demonstrates that GBJ can have significantly more power than popular alternatives such as GSEA or MAGMA while still protecting Type I error rate across various different pathway structures and also eliminating the need for computationally intensive genome-wide resampling.

Intuitively, GBJ and the goodness-of-fit methods owe their high power to two major factors. First, the structures of the test statistics allow for full incorporation of available GSA data when performing inference, in particular using the magnitude of each marginal summary statistic in the set as well as the joint SNP correlation structure. Secondly, these statistics are backed by strong theoretical results in simplified set-based settings, where they possesses asymptotic power guarantees. In finite samples, GBJ has been shown to provide
better performance than GHC.

In addition, we have provided a step-down inference procedure to mitigate the bias introduced through choice of a gene set definition file. Pathways that demonstrate strong associations based on a single gene are regularly identified as a serious problem^25^, ^52^, ^53^ and hinder important replication efforts. We show step-down inference can lessen these issues by highlighting the pathways that demonstrate effects over many genes, as opposed to pathways that rely on one or a few genes to drive their significance. Reporting only those findings that are still significant after the step-down procedure may help ensure that associations are replicable in studies with different pathway definitions.

One issue we have not discussed much is the philosophical difference between a self-contained test, such as the tests we have considered in this report, and a competitive test, such as certain variants of GSEA. In general, we recognize that both approaches possess unique strengths and weaknesses, and we believe both have their uses in GSA. Previous literature^6^ and the preceding work have demonstrated many of the advantages of self-contained tests, but there are certainly areas where a competitive analysis could provide additional benefits. In particular, a competitive test may have been able to provide more succinct lists of significant pathways by accounting for strong background signal present in the datasets we studied. However, we note that most studies will contain far less background signal, as the cohorts used in this paper are some of the largest ever assembled, and we have shown how GBJ is still able to provide useful inference even in highly polygenic settings. GBJ could also be recast as a competitive test using the gene permutation methods of other competitive strategies.

Another limiting factor for GBJ arises as a consequence of the data-intensive approach that affords it additional power. Very large gene sets containing over 1,500 SNPs can greatly slow down calculation of the test statistic and p-value, which can create difficulties analyzing the largest gene sets. While other tests may sacrifice large amounts of information by discarding more of the data, they can also produce results much more quickly as a consequence of utilizing fewer inputs. This issue can be alleviated by pruning or otherwise reducing the number of SNPs in a gene set so that GBJ still uses a large amount of information while running at an acceptable speed. Also, large amounts of data can cause issues with the default level of numerical precision in `R`, so that the current implementation of our software may not provide very precise p-values between 0 < *p* < 1 ⋅ 10^−12^. Still, 1 ⋅ 10^−12^ is generally a low enough significance level to account for multiple testing adjustments in GSA. GSEA, for example, can only provide a family-wise error rate as precise as 0.001 when testing a single pathway with its default of 1,000 permutations.

Generalized Berk-Jones represents a substantial departure from standard gene set analysis methods and offers distinct advantages over competing ideas, but there is still much room for future work. One possible extension would be a correction for background signal so that GBJ could provide an analytic p-value for the competitive null hypothesis. Computationally, it would be useful to develop algorithms that can calculate the statistic faster and with more precision so that larger gene sets can be tested quickly. Finally, it would be of interest to see how other set-based tests with similar asymptotic guarantees to Berk-Jones and Higher Criticism perform in the GSA paradigm. A number of such tests exist for independent summary statistics^32,54^ and could be modified to consider correlated data. These other methods may prove to provide even more finite sample power in the gene set analysis setting.

## Acknowledgments

We would like to acknowledge the Breast Cancer Association Consortium, the Psychiatric Genomics Consortium, and the GIANT Consortium for providing publicly available summary statistics.

## References

[1] Cantor RM, Lange K, Sinsheimer JS. Prioritizing GWAS results: a review of statistical methods and recommendations for their application. Am J Hum Genet. 2010;86(1):6–22.

[2] Locke AE, Kahali B, Berndt SI, Justice AE, Pers TH, Day FR, et al. Genetic studies of body mass index yield new insights for obesity biology. Nature. 2015;518(7538):197–206.

[3] Allen HL, Estrada K, Lettre G, Berndt SI, Weedon MN, Rivadeneira F, et al. Hundreds of variants clustered in genomic loci and biological pathways affect human height. Nature. 2010;467(7317):832–838.

[4] Nurnberger JI, Koller DL, Jung J, Edenberg HJ, Foroud T, Guella I, et al. Identification of pathways for bipolar disorder: a meta-analysis. JAMA Psychiatry. 2014;71(6):657–664.

[5] Visscher PM, Brown MA, McCarthy MI, Yang J. Five years of GWAS discovery. Am J Hum Genet. 2012;90(1):7–24.

[6] Fridley BL, Biernacka JM. Gene set analysis of SNP data: benefits, challenges, and future directions. Eur J Hum Genet. 2011;19(8):837–843.

[7] Pers TH. Gene set analysis for interpreting genetic studes. Hum Mol Genet. 2016;25:R133–R140.

[8] Yu K, Li Q, Bergen AW, Pfeiffer RM, Rosenberg PS, Caporaso N, et al. Pathway analysis by adaptive combination of p-values. Genet Epidemiol. 2009;33(8):700–709.

[9] Liu JZ, Mcrae AF, Nyholt DR, Medland SE, Wray NR, Brown KM, et al. A versatile gene-based test for genome-wide association studies. Am J Hum Genet. 2010;87(1):139–145.

[10] Mooney MA, Nigg JT, McWeeney SK, Wilmot B. Functional and genomic context in pathway analysis of GWAS data. Trends Genet. 2014;30(9):390–400.

[11] Wang K, Li M, Bucan M. Pathway-based approaches for analysis of genomewide association studies. Am J Hum Genet. 2007;81(6):1278–1283.

[12] Lips ES, Kooyman M, de Leeuw C, Posthuma D. JAG: a computational tool to evaluate the role of gene-sets in complex traits. Genes. 2015;6(2):238–251.

[13] Holmans P, Green EK, Pahwa JS, Ferreira MAR, Purcell SM, Sklar P, et al. Gene ontology analysis of GWA study data sets provides insights into the biology of bipolar disorder. Am J Hum Genet. 2009;85(1):13–24.

[14] Segre AV, DIAGRAM Consortium, MAGIC investigators, Groop L, Moothaa VK, Daly MJ, et al. Common inherited variation in mitochondrial genes is not enriched for associations with type 2 diabetes or related glycemic traits. PLoS Genet. 2010;6(8):e1001058.

[15] Lee PH, O’Dushlaine C, Thomas B, Purcell SM. INRICH: interval-based enrichment analysis for genome-wide association studies. Bioinformatics. 2012;28(13):1797–1799.

[16] Jia P, Zheng S, Long J, Zheng W, Zhao Z. dmGWAS: dense module searching for genome-wide association studies in proteinprotein interaction networks. Bioinformatics. 2011;27(1):95–102.

[17] O’Dushlaine C, Kenny E, Heron EA, Segurado R, Gill M, Morris DW, et al. The SNP ratio test: pathway analysis of genome-wide association datasets. Bioinformatics. 2009;25(20):2762–2763.

[18] de Leeuw CA, Mooij JM, Heskes T, Posthuma D. MAGMA: generalized gene-set analysis of GWAS data. PloS Comput Biol. 2015;11(4):e1004219.

[19] Gui H, Li M, Sham PC, Cherny SS. Comparisons of seven algorithms for pathway analysis using the WTCCC Crohns Disease dataset. Hum Genet. 2011;4(1):386.

[20] Jia P, Zhao Z. Network-assisted analysis to prioritize GWAS results: principles, methods and perspectives. Hum Genet. 2014;133(2):125.

[21] Evangelou M, Rendon A, Ouwehand WH, Wenisch L, Dudbridge F. Comparison of methods for competitive tests of pathway analysis. Bioinformatics. 2012;7(7):e41018.

[22] Jia P, Wang L, Meltzer HY, Zhao Z. Pathway-based analysis of GWAS datasets: effective but caution required. Int J Neuropsychopharmacol. 2011;14(4):567–572.

[23] Holmans P. Statistical methods for pathway analysis of genome-wide data for association with complex genetic traits. Adv Genet. 2010;72:141–179.

[24] Moskvina V, Schmidt KM, Vedernikov A, Owen MJ, Craddock N, Holmans P, et al. Permutation-based approaches do not adequately allow for linkage disequilibrium in gene-wide multi-locus association analysis. Eur J Hum Genet. 2012;20(8):890–896.

[25] Hong MG, Pawitan Y, Magnusson PK, Prince JA. Strategies and issues in the detection of pathway enrichment in genome-wide association studies. Hum Genet. 2009;126(2):289–301.

[26] Ramanan VK, Shen L, Moore JH, Saykin AJ. Pathway analysis of genomic data: concepts, methods, and prospects for future development. Trends Genet. 2012;28(7):323–332.

[27] Wu MC, Lin X. Prior biological knowledge-based approaches for the analysis of genome-wide expression profiles using gene sets and pathways. Stat Methods Med Res. 2009;18(6):577–593.

[28] de Leeuw CA, Neale BM, Heskes T, Posthuma D. The statistical properties of gene-set analysis. Nat Rev Genet. 2016;.

[29] Wang L, Jian P, Wolfinger RD, Chen X, Zhao Z. Gene set analysis of genome-wide association studies: methodological issues and perspectives. Genomics. 2011;98(1):1–8.

[30] Michailidou K, Lindstrom S, Dennis J, Beesley J, Hui S, Kar S, et al. Large-scale genetic association analysis identifies 65 new breast cancer susceptibility loci and predicts target genes. Nat Genet. 2017;551(7678):92.

[31] The Gene Ontology Consortium, Ashburner M, Ball CA, Blake JA, Botsein D, Butler H, et al. Gene Ontology: tool for the unification of biology. Nat Genet. 2000;25(1):25–29.

[32] Jager L, Wellner JA. Goodness-of-fit tests via phi-divergences. Ann Stat. 2007;35(5):2018–2053.

[33] Barnett I, Mukherjee R, Lin X. The Generalized Higher Criticism for testing SNP-set effects in genetic association studies. J Am Stat Assoc. 2017;112(517):64–76.

[34] Sun R, Lin X. Set-based tests for genetic association using the Generalized Berk-Jones statistic. arXiv, https://arxivorg/abs/171002469. 2017;.

[35] McCullagh P, Nelder JA. Generalized Linear Models. CRC Press; 1989.

[36] Chang CC, Chow CC, Tellier LC, Vattikuti S, Purcell SM, Lee JJ. Second-generation PLINK: rising to the challenge of larger and richer datasets. Gigascience. 2015;4(1):7.

[37] 1000 Genomes Project Consortium. A global reference for human genetic variation. Nature. 2015;526(7571):68–74.

[38] Fadista J, Manning AK, Florez JC, Groop L. The (in)famous GWAS P-value threshold revisited and updated for low-frequency variants. Eur J Hum Genet. 2016;24(8):1202–1205.

[39] Berk RH, Jones DH. Goodness-of-fit test statistics that dominate the Kolmogorov statistics. Z Wahrsch Verw Gebiete. 1979;47:47.

[40] Donoho D, Jin J. Higher criticism for detecting sparse heterogeneous mixtures. Ann Stat. 2004;32(3):962–994.

[41] Michailidou K, Beesley J, Lindstrom S, Canisius S, Dennis J, Lush M, et al. Genome-wide association analysis of more than 120,000 individuals identifies 15 new susceptibility loci for breast cancer. Nat Genet. 2015;47(4):373–380.

[42] Wood AR, Esko T, Yang J, Vedantam S, Pers TH, Gustafsson S, et al. Defining the role of common variation in the genomic and biological architecture of adult human height. Nat Genet. 2014;46(11):1173–1186.

[43] Schizophrenia Working Group of the Psychiatric Genomics Consortium. Biological insights from 108 schizophrenia-associated loci. Nature. 2014;511(7510):421–427.

[44] Gabregat A, Sidiropoulos K, Garapati P, Gillespie M, Hausmann K, Haw R, et al. The Reactome pathway knowledgebase. Nucleic Acids Res. 2015;44(D1):D481–D487.

[45] Thomas PD, Campbell MJ, Kejariwal A, Mi H, Karlak B, Daverman R, et al. Panther: a library of protein families and subfamilies indexed by function. Genome Res. 2003;13(9):2129–2141.

[46] Zhong H, Yang X, Kaplan LM, Molony C, Schadt EE. Integrating pathway analysis and genetics of gene expression for genome-wide association studies. Am J Hum Genet. 2010;86(4):581–591.

[47] Perry JRB, McCarthy MI, Hattersley AT, Zeggini E, the Wellcome Trust Case Control Consortium, Weedon MN, et al. Interrogating type 2 diabetes genome-wide association data using a biological pathway-based approach. Diabetes. 2009;58(6):286–292.

[48] Menashe I, Maeder D, Garcia-Closas M, Figueroa JD, Bhattacharjee S, Rotunno M, et al. Pathway analysis of breast cancer genome wide association study highlights three pathways and one canonical signaling cascade. Cancer Res. 2010;70(11):4453–4459.

[49] Aken BL, Ayling S, Barrell D, Clarke L, Curwen V, Fairley S, et al. The Ensembl gene annotation system. Database. 2016;.

[50] Johnstone RW, Frew AJ, Smyth MJ. The TRAIL apoptotic pathway in cancer onset, progression and therapy. Nat Rev Cancer. 2008;8(10):782–798.

[51] Manolio TA, Collins FS, Cox NJ, Goldstein DB, Hindorff LA, Hunter DJ, et al. Finding the missing heritability of complex diseases. Nature. 2009;461(7265):747–753.

[52] Elbers CC, van Eijk KR, Franke L, Mulder F, van der Schouw YY, Wijmenga C, et al. Using genome-wide pathway analysis to unravel the etiology of complex disease. Genet Epidemiol. 2009;33(5):419–431.

[53] Wang K, Li M, Hakonarson H. Analysing biological pathways in genome-wide association studies. Nat Rev Genet. 2010;11(12):843–854.

[54] Moscovich-Eiger A, Nadler B, Spiegelman C. On the exact Berk-Jones statistics and their p-value calculation. Electron J Stat. 2016;10(2):2329–2354.

